# Computational Modeling of the Biphasic Depletion of Ovarian Follicle Reserve and the Chemical Effects on Ovarian Aging

**DOI:** 10.1101/2025.03.27.645815

**Authors:** Sarahna A. Moyd, Shuo Xiao, Audrey J. Gaskins, Qiang Zhang

## Abstract

**Introduction:** Human ovaries begin development in utero. Through oogenesis, the numbers of oocytes and primordial follicles peak to a few million during fetal development, then decline to hundreds of thousands per ovary at birth. These primordial follicles do not regenerate and are thus regarded as the ovarian reserve. Over the life course, the reserve continues to deplete, due to atresia and activation, until menopause when about 1000 primordial follicles remain. Exposure to chemotherapy drugs and environmental pollutants can accelerate follicular depletion potentially leading to a greater risk of early menopause, primary ovarian insufficiency (POI), and infertility. Physiologically, the ovarian reserve is depleted in a seemingly biphasic pattern – a slow steady decline from birth to mid-30s, followed by a faster decline to menopause which typically occurs around age 50 years. While this depletion pattern has been described with empirical mathematical formulations, rarely is it modeled mechanistically. A mechanic model that can characterize the dynamics of follicular depletion throughout the life course will help researchers better understand and predict the impact of chemical exposures on ovarian aging.

**Methods:** Here we propose a minimal mechanistic model, which includes (1) a zero-order feedforward inhibition of primordial follicle activation by a local autocrine/paracrine inhibitory factor secreted by the primordial follicles, and (2) a high-gain feedback inhibition of primordial follicle activation by the anti-Müllerian hormone (AMH) secreted by the growing (primary, secondary, and early antral) follicles. The model is configured such that the two regulatory processes prevent primordial follicles from premature overactivation in early and late reproductive life stages, respectively. Two exposure scenarios - chemo-drugs/radiation and tobacco smoke - are presented to demonstrate predictive robustness and biological plausibility of chemically induced increases in cellular atresia.

**Results:** Our model recapitulates the biphasic depletion curve and predicts a constant supply of growing follicles through most of the active reproductive lifespan. This model predicts that the size of the initial primordial follicle pool plays the most significant role in determining menopausal age and suggests that unilateral ovariectomy may have a more attenuated effect than expected. Simulations of transient exposure to chemotherapy drugs provide an exposure example for promoting atresia of primordial and/or growing follicles and suggest exposure at earlier ages have greater impact on ovarian reserve and menopausal timing than exposure at later ages. Also, simulations of chronic chemical exposures suggest that chemicals which directly promote primordial follicle atresia are more damaging than chemicals directly promoting growing follicle atresia or inhibiting AMH, potentially leading to earlier age at menopause. A specific scenario of chronic exposure to cigarette smoke of various intensities was simulated to validate the prediction power of the model.

**Conclusions:** The ovary may have compensatory factors to extend reproductive age as long as possible amid insults that reduce the primordial follicle pool. The timing of these insults are likely an important variable. Future elaborations of such mechanistically based computational modeling with integration of in vitro toxicity testing data may help scaling efforts in predicting the implications of reproductive toxicants on ovarian aging.

## Introduction

### Ovarian depletion and aging phenomena

Infertility and impaired fecundity affect sizeable proportions of the U.S. population, across all demographics, family structures, and geographic locations. Approximately 15-25% of women between 15 to 44 years in the United States will experience infertility—the inability to get pregnant after one year of consistent, intentional attempts (Promotion 2022). Human ovaries begin development in utero. Through oogenesis, the number of oocytes and primordial follicles increase peaking at about 5-6,000,000 oocytes during fetal development, declining to about 100,000-500,000 per ovary at birth (Schenck et al. 2021; Wallace and Kelsey 2010). Primordial follicles, here referred to as nongrowing follicles (NGF), constitute the ovarian reserve. These follicles steadily decline across a woman’s reproductive lifespan. To maintain the most viable pool of NGF from birth until menopause, the vast majority remain inactive, or dormant during early stages of development. After the onset of puberty via paracrine and autocrine signaling, the NGF pool collectively optimizes each follicle’s longevity until atresia—the ultimate and normal fate of >99% of the NGFs (Zhao et al. 2021). At menopause, typically fewer than 1,000 NGF remain in both ovaries combined (Broekmans, Soules, and Fauser 2009; Faddy et al. 1992). The depletion of NGFs result from both atresia and their activation to growing follicles (GF). GFs initially undergo a lengthy gonadotrophin-independent folliculogenesis phase where primordial follicles slowly develop into primary, secondary, and early antral follicles, and then a small fraction of GFs undergo gonadotrophin-dependent folliculogenesis where one dominant antral follicle is selected for ovulation (Zeleznik 2004). Approximately 30 large antral follicles enter the gonadotropin-dependent phase at the beginning of each cycle, then a single is selected as its growth outpaces others. The unselected follicles undergo atresia.

It is widely accepted that ovarian follicles do not regenerate, therefore any harms including acute or chronic exposure to environmental chemicals, particularly those classified as endocrine disrupting chemicals (EDCs) and genotoxic compounds that have been demonstrated to cause premature depletion of the reserve (’<dissertation_dingning_1.pdf>’ ; Hirshfield 1994a; Wang et al. 2019b; Xu et al. 2020), are important to identify for mitigation and protection. The ovary exhibits unique aging mechanisms that outpace cellular aging of other somatic tissues, making it particularly vulnerable to cellular dysregulation (Olsen et al, 2020) Premature reductions to the reserve may have irreversible reproductive consequences, such as primary ovarian insufficiency (POI) also seen in the literature as premature ovarian failure (POF), or premature menopause (Ding et al. 2020; Podfigurna-Stopa et al. 2016). POI is a disorder characterized by a low-for-age oocyte count and decreased AMH levels, a hormone that is secreted by the healthy, developing secondary, preantral, and early antral follicles as part of the GF pool (Pastore et al. 2018; Gynecologists 2014; Ding et al. 2020). Diminished ovarian reserve (DOR) is a clinically complex diagnosis as it may encompass women with an abnormal ovarian reserve test, a previous poor response to ovarian stimulation, and/or advanced maternal age. For example, women aged 40+ years with DOR may not have underlying pathology, but simply fewer remaining oocytes given their age; however, women <35 years may be classified as DOR due to low AMH/AFC levels but still have high-quality oocytes thereby not necessarily reducing probability of conception (Pastore, et al 2018). Premature menopause is defined as the occurrence of natural menopause (e.g. primordial follicle count less than 1,000) prior to 40 years of age (Podfigurna-Stopa et al., 2016; DeVos et al., 2010). Age is not only the most important predictor of ovarian reserve and fertility outcomes, but with age also comes declined oocyte quality via accumulated environmental exposures and reduced capacity to overcome exogenous insults resulting in decreased oocyte viability, further contributing to infertility (Domínguez et al., 2016; Masayuki Shimada & Yasuhisa Yamashita, 2011). Considerable knowledge gaps and mechanistic understanding of oocyte development, maturation, and ovarian aging remain, which this study aims to address.

### Biphasic-like depletion of NGF

Previous studies report quantitative statistical models of the age-associated follicle decline, suggesting a biphasic decay — two characteristically distinct linear phases on the log scale of NGF with respect to age, a slow, steady phase followed by a faster decay (Faddy et al. 1992; Broekmans, Soules, and Fauser 2009; Wallace and Kelsey 2010; Kelsey et al. 2011b). The rate of follicular decline appears to be ∼0.097 of the primordial follicle pool/year when at least ∼25,000 NGF remain, then shifts to ∼0.237 of the primordial follicle pool/year, a greater than 2-fold increase (Faddy and Gosden 1996). The first phase can be described as the ovarian reserve decreasing linearly by approximately 50% from birth to puberty, then to about 10% by age 30 (Wallace and Kelsey 2010; Upson et al 2019). The second phase takes over usually in a woman’s mid-30s with NGF depletion accelerating until menopause at around age 50 years (Broekmans, Soules, and Fauser 2009). There is debate regarding if this behavior is truly biphasic—whether the two states truly represent a drastic or abrupt rate of follicular atresia (Leidy et al., 1998), however it appears more reasonable that with age also comes a breakdown in the neuroendocrine feedback/feedforward mechanisms essential to inhibiting overactivation of primordial follicle early in life. It is estimated when ∼100,000 NGF remain, menopause is expected to occur in ∼21.5-26.5 years, and when ∼10,000 NGF remain, menopause is expected to occur in ∼5-10 years (Faddy and Gosden 1996).

### Uncertainty in mathematically describing ovarian follicle depletion

Early studies report the first phase decays exponentially while later studies, particularly by Wallace and colleagues, provide evidence of the first phase decay behaving more linearly (Broekmans et al., 2009b; Faddy et al., 1992; Hansen et al., 2008; Seifer et al., 2011; Wallace & Kelsey, 2010). Further, Wallace and colleagues assert NGF deplete at a rate of about 3% per year from birth to age 30. Combining evidence from various studies, Park and colleagues also report linear depletion from age 15 to late 20s (Park et al., 2022). This trend is also seen in mice with a linear decline observed between the 2^nd^ and 3^rd^ months of life (Park, Oh et al. 2022). The later studies are typically deemed more reliable due to the greater availability of human data as Wallace and colleagues obtained follicle counts from a few hundred women compared to the much smaller sample sizes of earlier studies. Furthermore, early studies characterize the declining follicle counts with age via two exponential decay equations for both phases, with the exponential equation describing the second phase having a higher decay rate constant (Kelsey et al. 2011b; Faddy et al. 1992; Broekmans, Soules, and Fauser 2009; Iyer et al. 2019; Lie Fong et al. 2012). As a result, two curves of different slopes are combined to describe the relationship between log values of primordial follicles and age in years. Later studies including those by Wallace, Kelsey, & Wright *et. al* started implementing smooth functions to characterize the depletion. For example, Wallace used second-order power function to describe the entire biphasic depletion (Wallace and Kelsey 2010), and Hansen used a third-order function to describe this relationship with the fitted relationship between log values of primordial follicles and age appearing concave down (Hansen et al. 2008). Despite a dearth of mechanistic descriptions, these seminal mathematical models have aided in gleaning insight into a phenomenon not readily observable.

### Conserved Mechanism for Constant supply of GF

It is biologically reasonable to state the number of GF recruited during reproductive age remains relatively constant in each cycle as opposed to decreasing proportionally to the size of the declining pool of primordial follicles. The number of early GF exhibits decline within a much narrower fold-range than NGF with age (Gougeon, Ecochard, and Thalabard 1994). A constant number of follicles entering the growing phase over time despite declining reserve has also been observed in rats (Hirshfield 1994a). Similar patterns are observed in mice: while primordial follicles decrease by nearly 17-fold from 25 days to 13 months of age, the small growing follicles only decreased by 50% (Durlinger et al., 1999). The increasing fraction of early GF and decreasing fraction of NGF with age suggests regulated activation of NGF over the reproductive lifespan (Gougeon & Chainy, 1987). If the depletion of the NGF is caused predominantly by activation as opposed to atresia, then a linear decay in the first phase predicts a constant supply of GF during the active reproductive ages. However, it appears that atresia may dominate NGF depletion during early reproductive age, followed by a switch at around age 30 years where activation dominates depletion (Gougeon, Ecochard, and Thalabard 1994). Should atresia truly dominate depletion, regulatory mechanisms must exist to ensure a constant supply of GF. Without such mechanisms, the activation rate is likely to be proportional to the number of remaining NGF, and the supply of GF should not be expected to remain constant as the reserve depletes. A constant pipeline of growing follicles over the reproductive life span also supports the age-associated linear decline until around 37 years when exponential depletion becomes more evident (Wallace & Kelsey, 2010). This observed dynamic of regulated recruitment and depletion appears to be controlled by key ovarian hormones via tiered inhibition: feedforward by NGF via an autocrine inhibitory factor, then feedback inhibition by GF-secreted Anti-Müllerian Hormone (AMH) (Hansen et al. 2008).

### Mechanism of regulated follicle recruitment and depletion

AMH is a member of the transforming growth factor β (TGF-β) family (Anckaert et al. 2012). In the ovary, AMH is only secreted by primary, secondary, and early antral follicles (Dumont et al., 2015). Intraovarian AMH plays a crucial role in regulating folliculogenesis as it inhibits both (1) the gonadotrophin-independent initial recruitment of NGF to the GF pool, and (2) the FSH-dependent cyclic recruitment of small antral follicles into waves of cohort available for dominant follicle selection in each menstrual cycle (Baerwald, 2009; Baerwald et al., 2012; Mcgee & Hsueh, 2000). Serum AMH, which mirrors intraovarian AMH, increases by 2-2.5 fold from birth to its peak between ages 18-20 (Kelsey et al. 2011b; Lie Fong et al. 2012; Hagen et al. 2010), then plateaus through early 20s before decaying with the depletion of ovarian reserve. By age 35, AMH has decreased to approximately half its peak value, then only ∼5 years later it reduces by half again; by menopause AMH has diminished completely (Kelsey et al. 2011b; Lie Fong et al. 2012; Hagen et al. 2010; Iyer et al. 2019; Seifer, Baker, and Leader 2011). Therefore, serum AMH can be regarded as a strong biomarker for predicting ovarian aging, however, due to high inter-individual variability, there are still lingering challenges to solely relying on serum AMH as a predictor of ovarian reserve, oocyte quality, and response to ovarian stimulation (Broekmans et al., 2009a; Lie Fong et al., 2012; Seifer et al., 2011).

Another, elusive, poorly characterized regulatory signal that is hypothesized to play an important role in NGF depletion is NGF inhibitory factor, herein referred to as InhF. This NGF-associated factor is suggested to exist and have a role in growth inhibition prior to puberty when AMH levels are low. This inhibitory indicator has been referenced in a number of studies and described to work via autoregulation and paracrine signaling (Da Silva-Buttkus et al. 2009; Hirshfield 1994a; Schenck et al. 2021). An early study showed that the fraction of growing follicles is inversely correlated with the number of primordial follicles, suggesting a self-inhibitory mechanism is in place, although whether it is mechanical or autocrine in nature is unclear (Hirshfield 1994b). In the mouse ovary from birth to day 12, primordial follicles with more primordial follicles as neighbors are less likely to be activated to growth (Da Silva-Buttkus et al. 2009). The non-random, clustered distribution of primordial follicles observed within the ovary at older age strongly suggests these follicles secrete an autocrine/paracrine factor to inhibit their own growth activation. The inhibition may be synergistic because as the number of clustered follicles within a short spatial distance (10 microns) increases, i.e. increased density of NGF clusters, a greater than additive inhibition is observed (Schenck et al. 2021). It has also been suggested that as density decreases, the remaining primordial follicles sustain the clusters. Such density-dependent autocrine-paracrine inhibition acts to preferentially conserve the pool of primordial clusters of higher density over the life span such that the reserve is not depleted uniformly in the ovarian cortex. Additionally, NGF closer to the ovarian medulla tend to be depleted earlier than those closer to the ovarian surface, which is consistent with the InhF hypothesis because the local concentration of InhF at the cortex/medulla border is expected to be the lowest (Schenck et al. 2021). Taken together, these studies further reiterate the truly intricate nature of the ovary and its self-preserving system of network communication.

### Hypothesis

We hypothesize the coordinated action of the InhF-mediated incoherent feedforward and AMH-mediated negative feedback maintains a constant supply of activated GF through reproductive years (Fig. 1). Specifically, through incoherent feedforward mediated by InhF, follicle activation is a zero-order process, i.e., independent of the number of remaining NGF. Secondly, as the reserve gradually depletes and the inhibition exerted by the InhF wanes due to lower density, AMH-mediated negative feedback dominates to continuously maintain a relatively constant supply of GF. After mid-30s, the decline in AMH leads to negative feedback breakdown, resulting in decreased inhibition of NGF recruitment and thus accelerated depletion through reproductive senescence (Da Silva-Buttkus et al., 2009). The phenomenon of follicle activation and depletion is scarcely described mathematically based on biology, which has prevented the development of a single mechanistically based model until now. Here, based on our hypothesis, we 1) present a computational model incorporating current mechanistic knowledge of follicle activation and depletion to predict ovarian aging, and 2) test hypotheses with simulated populations on the effects chemical exposure on ovarian aging.

**Figure 1.**
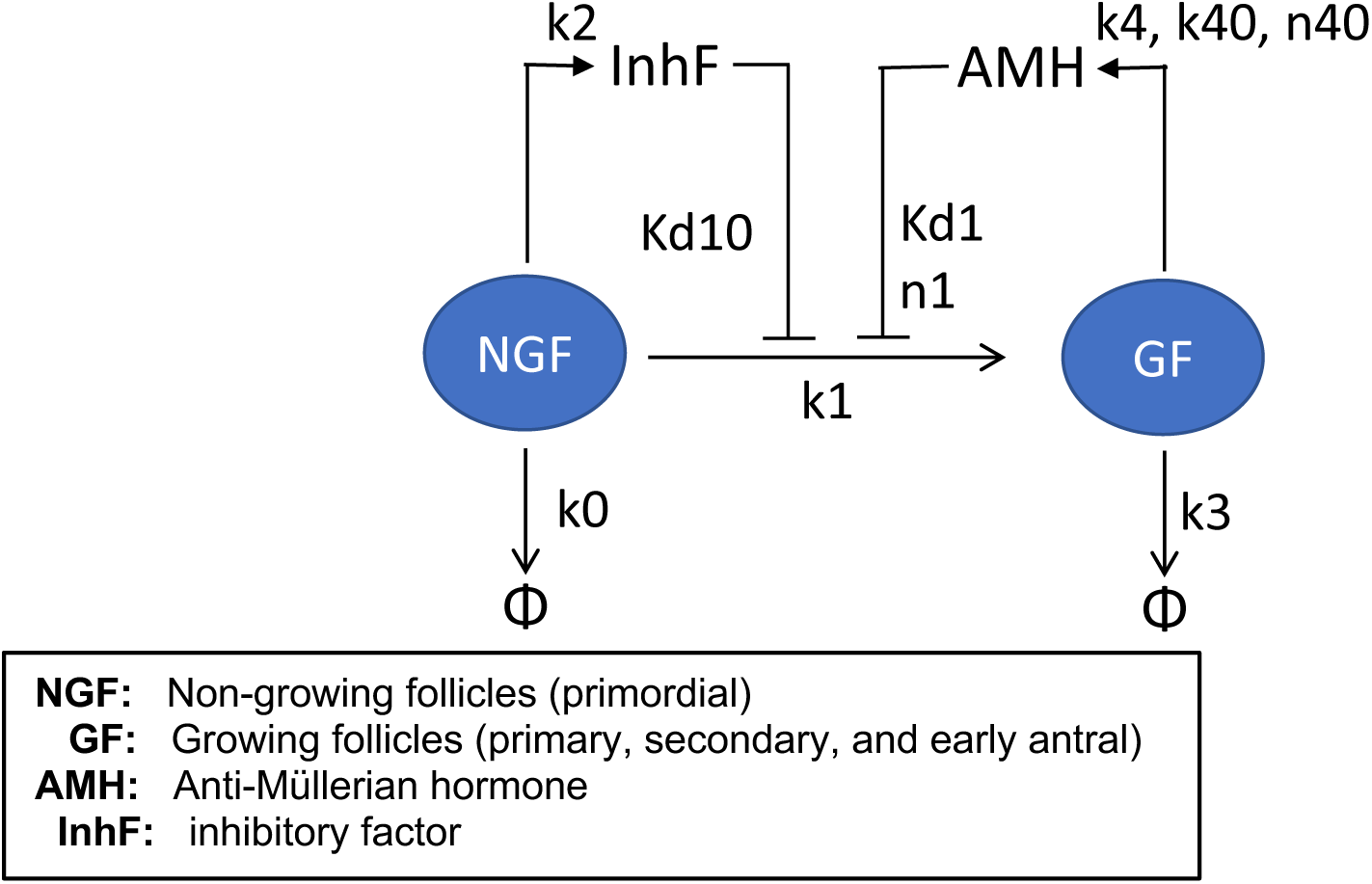
Model structure of NGF activation and depletion.

## Methods

### Model structure and assumptions

Figure 2.1 presents the structure of a minimal model of NGF depletion through atresia and activation to GF. There are 2 intraovarian local factors: InhF secreted by NGF and AMH secreted by GF. Since the time scales of InhF and AMH turnover are likely negligibly short relative to the reproductive lifespan, these 2 factors are assumed to be in quasi-steady state and change proportionally with the numbers of NGF and GF, respectively. AMH level is also age-dependent and is described by a saturable Hill function such that AMH increases from birth through puberty. The following two equations describe the level of InhF and AMH:

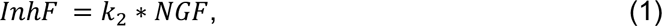

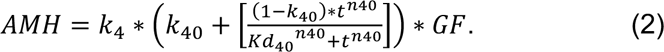

*k*_2_ is the scaling constant for InhF with NGF, *k*_4_ is the scaling constant for AMH with GF; *k*_40_, *Kd*_40_ and *n*_40_ are parameters describing the age-dependence of AMH.

NGF has 2 fates: (1) atresia which is assumed to be first-order, or (2) activation to GF. Activation is controlled by feedforward inhibition by InhF and feedback inhibition by AMH. The ordinary differential equations (ODEs) describing the time dependent NGF and GF are:

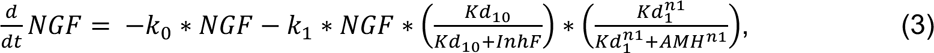

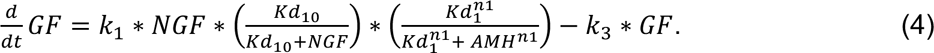

*k*_0_ is the first-order rate constant for NGF atresia, *k*_1_ is the activation rate constant for NGF to GF, *Kd*_10_ is the Michaelis constant for InhF inhibition of NGF activation; *Kd*_1_ and *n*1 are the affinity constant and Hill coefficient, respectively, for the AMH-mediated inhibition of activation of NGF; and *k*_3_ is the first-order rate constant for GF atresia.

### Human data and model parameterization and optimization

Early work compiled 8 quantitative histological studies of NGF, totaling 315 human samples. Eleven of these samples were from fetuses and infants (BLOCK, 1953) and only provide an initial NGF count. All others provided counts of NGF and GF (BLOCK, 1952, 1953; Gougeon et al., 1994; Richardson et al., 1987). Some samples were obtained from gynecological surgery i.e. unilateral oophorectomy/oophorectomy (Post et al., 2008), elective termination (Bendsen et al., 2006), fetal, neonate, infant death (Forabosco & Sforza, 2007), and others postmortem from unexpected child or adult death. Data from these human samples were used to generate distributions of NGF counts with respect to ages. A subset of data from the Environment and Reproductive Health Study (EARTH) prospective preconception cohort study were used to generate distributions of GF counts with respect to age (Messerlian et al. 2018)

Computational modeling was executed in MATLAB and ODEs were solved with the ode15s solver (*MathWorks Inc.*, 2023). Top sensitive model parameters were optimized using MATLAB’s fmincon function which finds the minimum solution to the specified objective, or cost, function within specified upper and lower bound constraints. Our cost function minimizes the squared difference between model prediction and human NGF count data described above.

### Sensitivity analysis

Local sensitivity analyses were conducted to describe the influence of model parameters on endpoints including menopause age or NGF count at a given age. This was performed by increasing and decreasing one parameter at a time by 1% from the default value and calculating the relative sensitivity coefficient for each endpoint by averaging the normalized percentage changes of the endpoints in both directions.

### Monte Carlo (MC) simulation of a virtual population

To simulate a virtual woman population, the 4 most sensitive parameters were selected for MC simulation. The initial NGF count follows a lognormal distribution as provided in Johnson et al. (Johnson et al., 2022). The remaining parameters k0, k1, k2, k3, k4, k40, Kd1, Kd10, Kd40, Kd41, n1, n40, n41 also follow respective lognormal distributions with the means equal to optimized default values and variances as listed in Table 1. Simultaneous random sampling was performed for the 4 parameters from the above-described distributions 1000 times and the values were used as input to the ODE model to simulate a population size of 1000 women. In our simulation, the model assumes two independent ovaries for each individual with both ovaries having identical parameter values. When remaining NGF per ovary reaches 500 (1000 NGF total across both ovaries), the individual was classified as having achieved the outcome of menopause. This threshold of 1000 across both ovaries for menopause is also used in prior studies (Faddy and Gosden 1996).

**Table 1:** Model parameter Values.

| Param | Description | Value |
| --- | --- | --- |
| init.NGF | NGF per ovary at birth | 0.325E6 |
| init.GF | GF per ovary at birth; the NGF:GF ratio is 30:1 | 1475 |
| k0 | rate constant for NGF atresia | 0.35152 |
| k1 | maximum rate constant for NGF activation | 0.23 |
| k2 | scaling factor for InhF production by NGF | 1 |
| k3 | rate constant for GF atresia | 4 |
| k4 | scaling factor for AMH production by GF | 10 |
| k40 | Age-dependent scaling factor for AMH production by GF | 0.2 |
| Kd1 | Effective affinity constant for AMH inhibition of | 6650 |
| Kd10 | Effective affinity constant for InhF inhibition of NGF activation | 32900 |
| Kd40 | Effective affinity constant for age-dependent AMH production by GF | 11 |
| Kd41 | Effective affinity constant for enforced age-dependent AMH production | 32 |
| n1 | Hill-coefficient for InhF inhibition of NGF activation | 4 |
| n40 | Hill-coefficient for age-dependent AMH production by GF | 1.5 |
| n41 | Hill-coefficient for enforced age-dependent AMH production | 7 |
| k_zero_order | rate constant for zero-order NGF depletion | $(300000-36000)/30$ |
| k_first_order | rate constant for first-order NGF depletion | 0.07067 |

### Simulation of chemical effects

Two exposure scenarios were executed to obtain ages of menopause that can be compared to literature. Both chemo-drugs/radiation used in cancer therapy and tobacco smoke can induce atresia due to genotoxicity or other cellular stress-related apoptosis leading to POI (Andersen et al., 2019; Xu et al., 2020). The effects of ovarian toxic chemicals that may accelerate NGF depletion were simulated according to potential mechanisms, where the atresia of NGF or GF or the secretion of InhF or AMH may be impacted singly or in combination. Some chemo-drugs, such as doxorubicin and cyclophosphamide, are postulated to increase both atresia of NGF and apoptosis of GF (Kalich-Philosoph et al. 2013; Wang et al. 2019a). This would involve increasing the values of model parameters k0 and/or k3 if altered atresia is involved, or decreasing the values of k2 or k4 if inhibition of InhF and AMH secretion is involved. Effects can also be nonlinear. For instance, when NGF is reduced due to increased atresia, InhF is also reduced which will accelerate NGF activation and thus depletion. Similarly, we tested a dose-effect presuming 20 years of smoking, which is the average duration among chronic smokers and spans the life course from after puberty (age 15) through a woman’s mid 30s. Tobacco smoke exposure reduces GF which in turn reduces AMH level, thereby accelerating NGF activation and depletion.

## Results

Given the uncertainty of the NGF depletion curve, especially for the first phase as described above, two reference curves were generated to represent either exponential or linear NGF depletion (Figure 2). A benchmark age of 30 years was used as the anchor point, at which NGF has depleted to 12% of the initial NGF count at birth based on prior studies (Mattison, 1989; Wallace & Kelsey, 2010). The two reference curves are presented in Fig. 2A and 2B for the linear and log scale of NGF respectively. Exponential decay represents unregulated NGF depletion where each follicle is removed from an ovary (via activation or atresia) independently, with a fixed first-order rate constant. With exponential decay, the half-life of NGF is calculated to exceed 10 years. In contrast, linear decay represents highly regulated NGF depletion where a constant number of follicles are removed per unit time, i.e., zero-order decay. Note that when displayed on logY scale the exponential decay becomes a straight line (Fig 2B, red line), whereas the linear decay appears to be biphasic – initially linear followed by accelerated decay by mid-30s (Fig 2B, blue line). From birth to age 30 years, the exponential decay model depletes NGF faster than the linear decay model then after age 30 the depletion rate of the exponential decay model decreases compared to the linear decay model. Biologically, NGF depletion is expected to follow a trend that is between the two reference cases.

**Figure 2.**
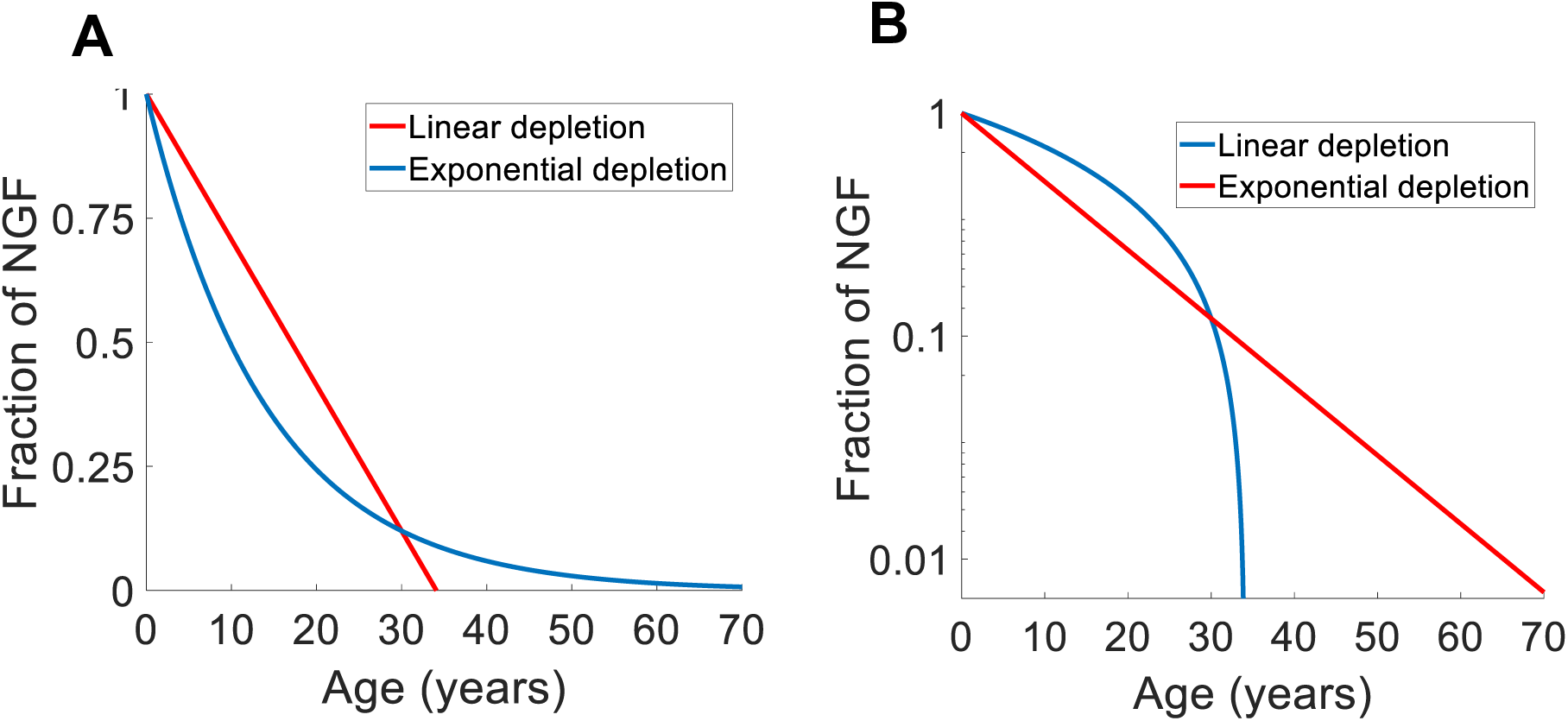
Reference depletion curves on (A) linear and (B) log scale.

### Model-simulated average NGF depletion

Using the human data and parameter optimization process as described in Methods, the model produced an average NGF depletion curve that fit the population data with R^2^=0.67 (Fig. 3A). The NGF depletion curve (blue) is biphasic with an initial slow phase from birth to mid-30s followed by a faster phase in late reproductive years. In comparison, while GF are also biphasic (pink), it has a much smaller slope in the initial phase, indicating that despite NGF dropping by >90% by women’s mid-30s, a steady GF count is maintained, declining more slowly percentwise. Interestingly, the model simulating average NGF curve is in between the exponential and linear decay reference curves (Fig. 3B). The NGF atresia and activation rates per year are shown in Fig. 3C where initially atresia is dominant and gradually yields to activation by the age of 30 years. The intraovarian InhF and AMH levels are shown in Fig. 3D, where InhF has the same profile as NGF, while in contrast AMH showed a typical bell-shaped profile that peaks around age 20 years, as expected.

**Figure 3.**
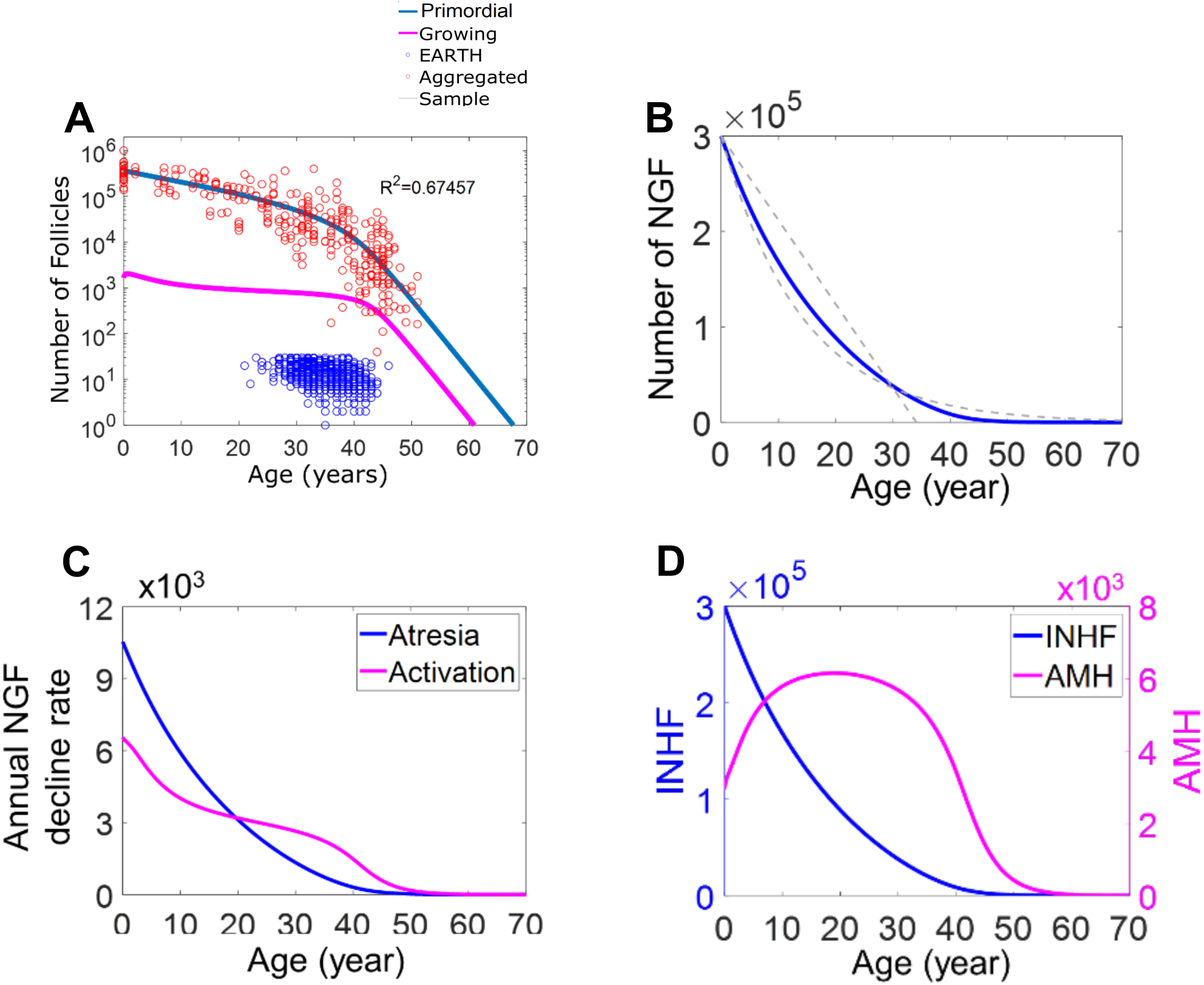
Model simulated average NGF depletion. (**A**) Model simulated NGF (primordial) and GF. Aggregated data of NGF and GF (red circles) are from n=315 human study participants (Wallace and Kelsey 2010b). Wallace and Kelsey 2010, antral follicle count is from EARTH study (Messerlian et al. 2018). **(B)** The model simulated NGF curve is in between the exponential and linear decay reference curves. **(C)** NGF atresia and activation rates (number of follicles per year) **(D)** Intraovarian INHF and AMH levels (arbitrary unit).

### Sensitivity analysis

Tornado plots were used to visualize parameter sensitivity (Fig. 4). Local sensitivity analysis showed that the most important parameter in predicting menopause age is the initial NGF count, followed by k0, k1, and k4 (Fig. 4A). A similar set of parameters were also identified as being the most sensitive to affecting the NGF remaining at ages 35 and 50 years (Fig. 4C and 4D). For the NGF remaining 1 year after birth, the initial NGF is the only influential parameter (Fig. 4B).

**Figure 4.**
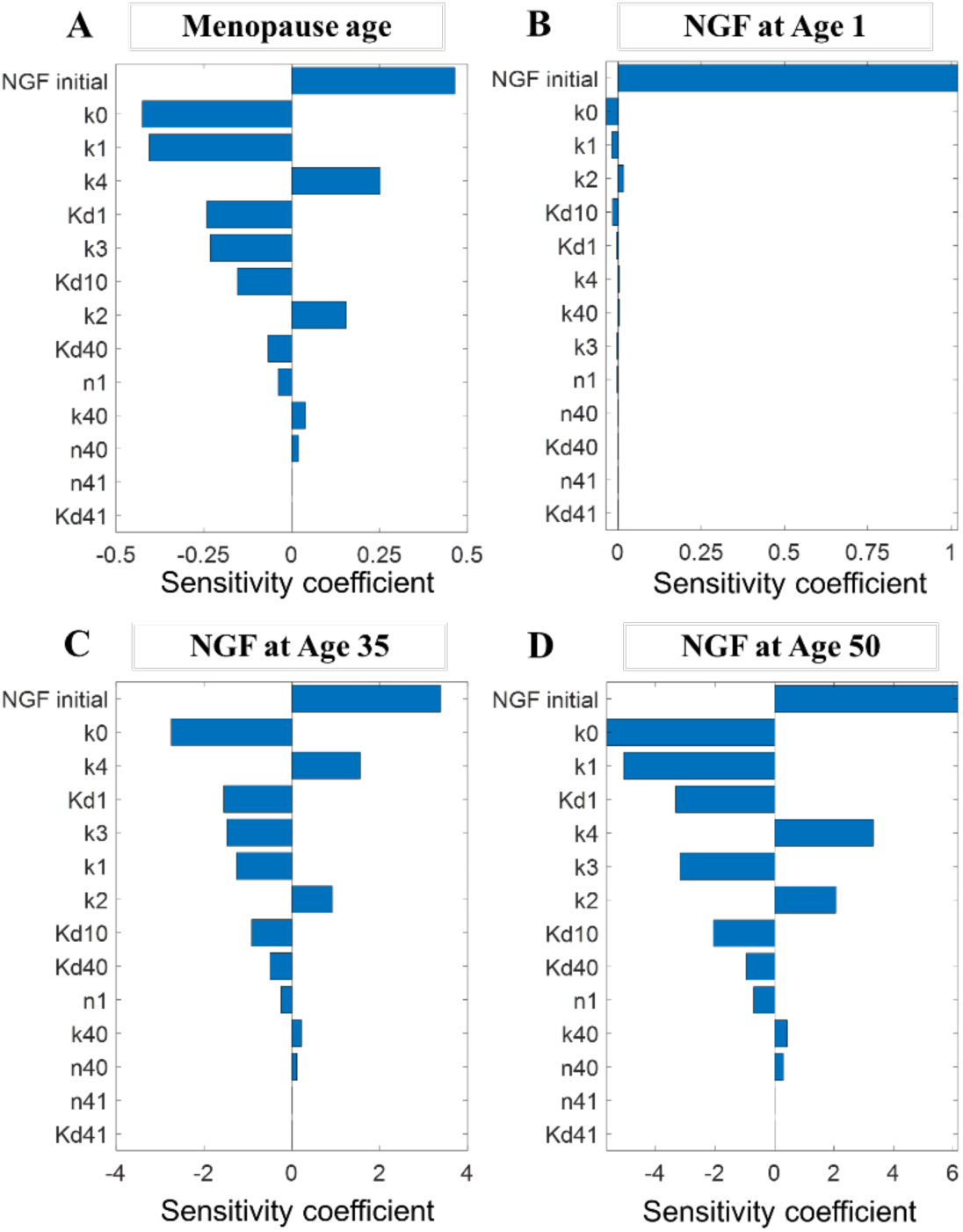
Tornado plots of local parameter sensitivity. Model parameter importance by (A) menopausal age predicted by model, (B-D) ages across the lifespan of age 1, age 35, and age 50.

### MC population simulation

MC simulation shows that the human population NGF depletion data can be adequately described by an ensemble of 1000 individuals (Fig. 5A). When using 500 NGF remaining per ovary as the cutoff threshold for predicting menopause age, the predicted menopause age distribution has a mean ± sd of 50.3 ± 4.3 years and a median of 50.1 years (Fig. 5B). To validate this distribution, the model-predicted cumulative probability was overlayed with 3 cumulative probability datasets reported for menopause age in different populations (Kawai, 2012; Pokoradi et al., 2011; Weinstein et al., 2003). The results showed that the model prediction falls within the range of the cumulative distributions in the literature (Fig. 5C).

**Figure 5.**
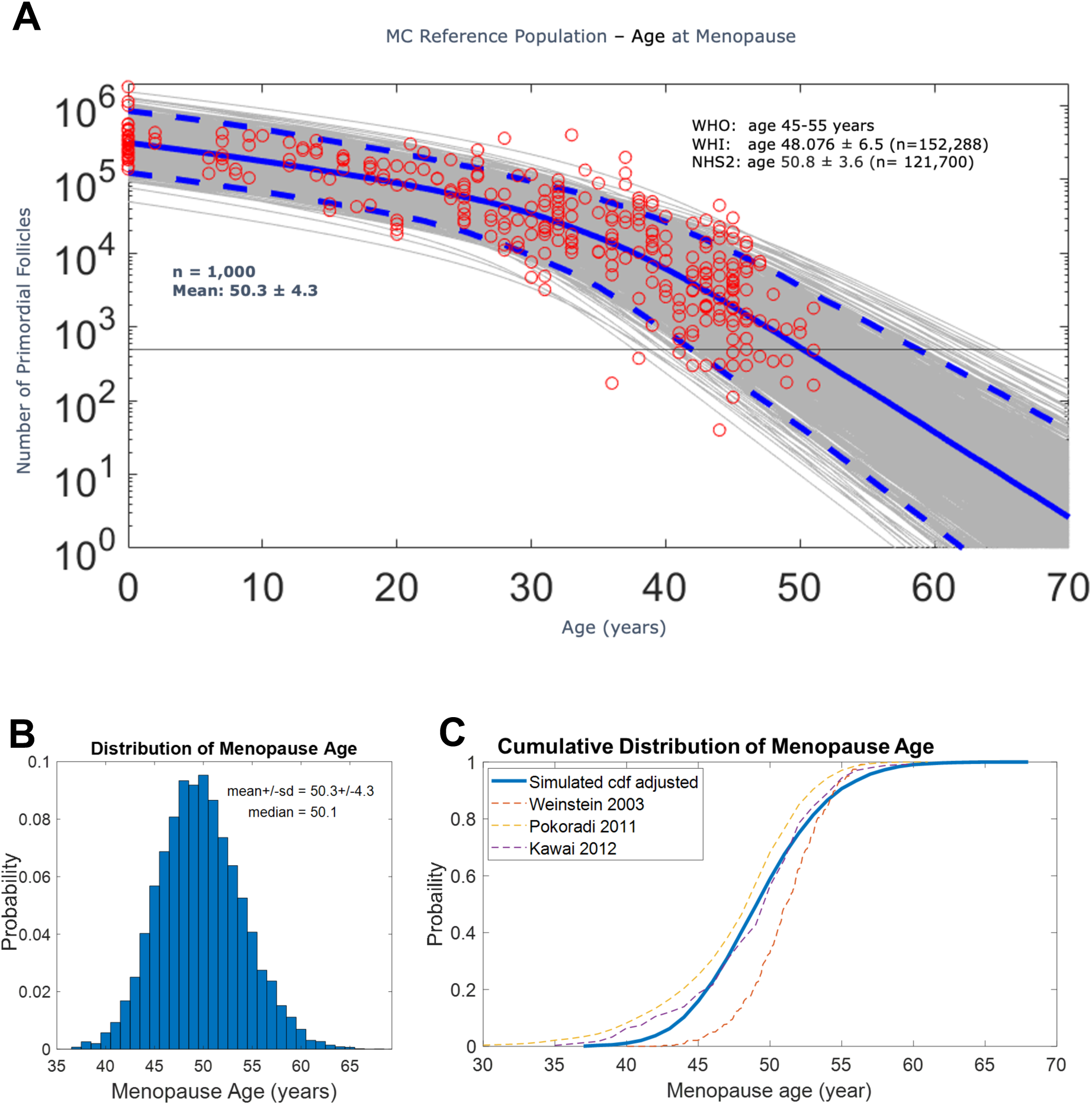
MC simulation of NGF depletion. (**A**) 1000 individual MC simulations (grey lines) with mean (solid blue) and 95% reference range (dash blue) indicated. Red circles are n=315 human samples that provide counts of NGF and GF (Kelsey et al. 2011a). Horizontal grey line: 500 NGF per ovary threshold for menopause.Age at menopause reported by the World Health Organization (WHO), the Women’s Health Initiave (WHI), and the Nurses Health Study II are indicated as well (Hu et al. 1999). **(B)** Model-predicted menopause age distribution with mean, sd and meadian as indicated. **(C)** Model-predicted cumulative distribution of menopause age overlayed with literature data as indicated.

### Simulated generic chemical effects

We first simulated a scenario for continuous, lifetime environmental chemical exposure. Assuming the simulated chemical can increase the atresia rate of NGF by 50% at a given exposure level, the simulation showed that although the NGF depletion accelerates such that the mean menopause age lowers, the advancement is nonlinear. The mean menopause age is approximately 8 years earlier (Fig. 6A), and not proportional to the percentage increase of the NGF atresia rate. This result indicates that the feedforward and feedback mechanisms implemented in the model allow the NGF to deplete slower than an unregulated system would do. We next simulated scenarios of short-term acute exposure to chemo-drugs which reduce NGF through atresia. Simulation is parameterized for a one-year exposure to a chemo-drug that increases k0 by 10-fold at age 10. The model predicts an earlier mean menopause age of 46 years old (Fig. 6B), while if the same chemotherapy occurs at age 20, the impact is smaller whereas the mean menopause age is 47.3 years (Fig. 6C).

**Fig. 6.**
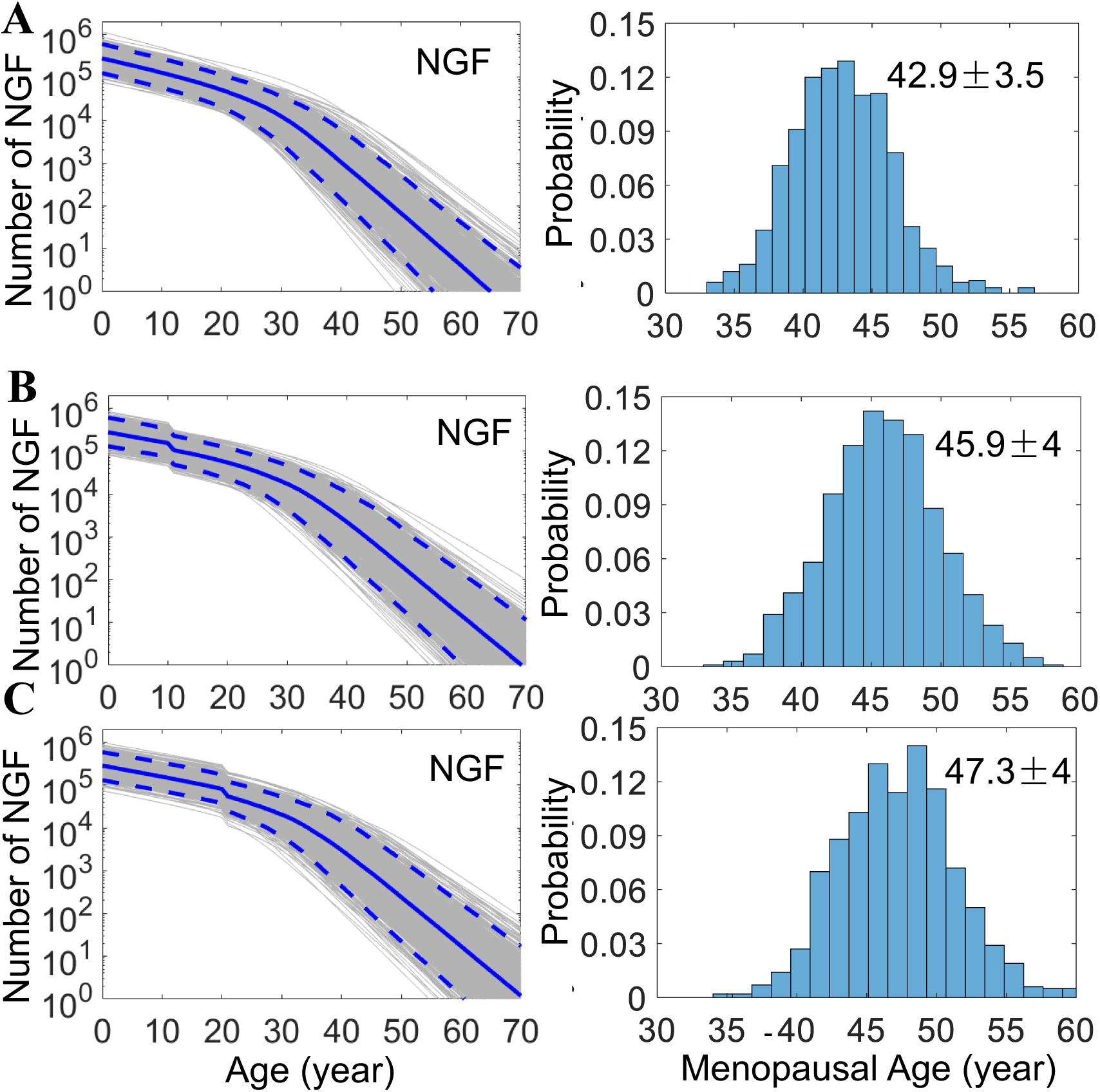
Simulation results for chemical effects. (**A**) Accelerated NGF depletion and advanced menopausal age for life-long exposure to a chemical that increases primordial follicle atresia rate (k0) by 50%. Accelerated NGF depletion and advanced menopausal age for one-year exposure at **(B)** age 10, and **(C)** age 20 respectively to a chemo-drug that increases k0 by 10 fold. Mean±SD is indicated for menopausal age.

### Effects of cigarette smoke exposure

In this sub analysis, we simulated the cigarette smoke exposure scenario by assuming individuals started smoking at age 15 and quit at age 35. The impact of smoking on atresia of primordial follicles depends on the number of cigarettes smoke per day. It has been estimated that for heavy smokers of >20 cigarettes per day the atresia rate can double (Mattison, 1989). We therefore implemented an increase in k0 from 1.2-2-fold to simulate these different smoking scenarios. A k0 of 1.2 was used for nonsmokers to approximate second-hand smoking or other similar background exposure scenarios. The MC simulations showed that as the degree of cigarette smoking increases, the cumulative probability of menopause age shifts to younger ages (Fig. 7A). There appears to be a linear inverse relationship between the degree of smoking and the mean menopause age (Fig. 7B). For heavy smokers, the mean menopause age is accelerated to 43.9 years (Fig. 7B).

**Fig. 7.**
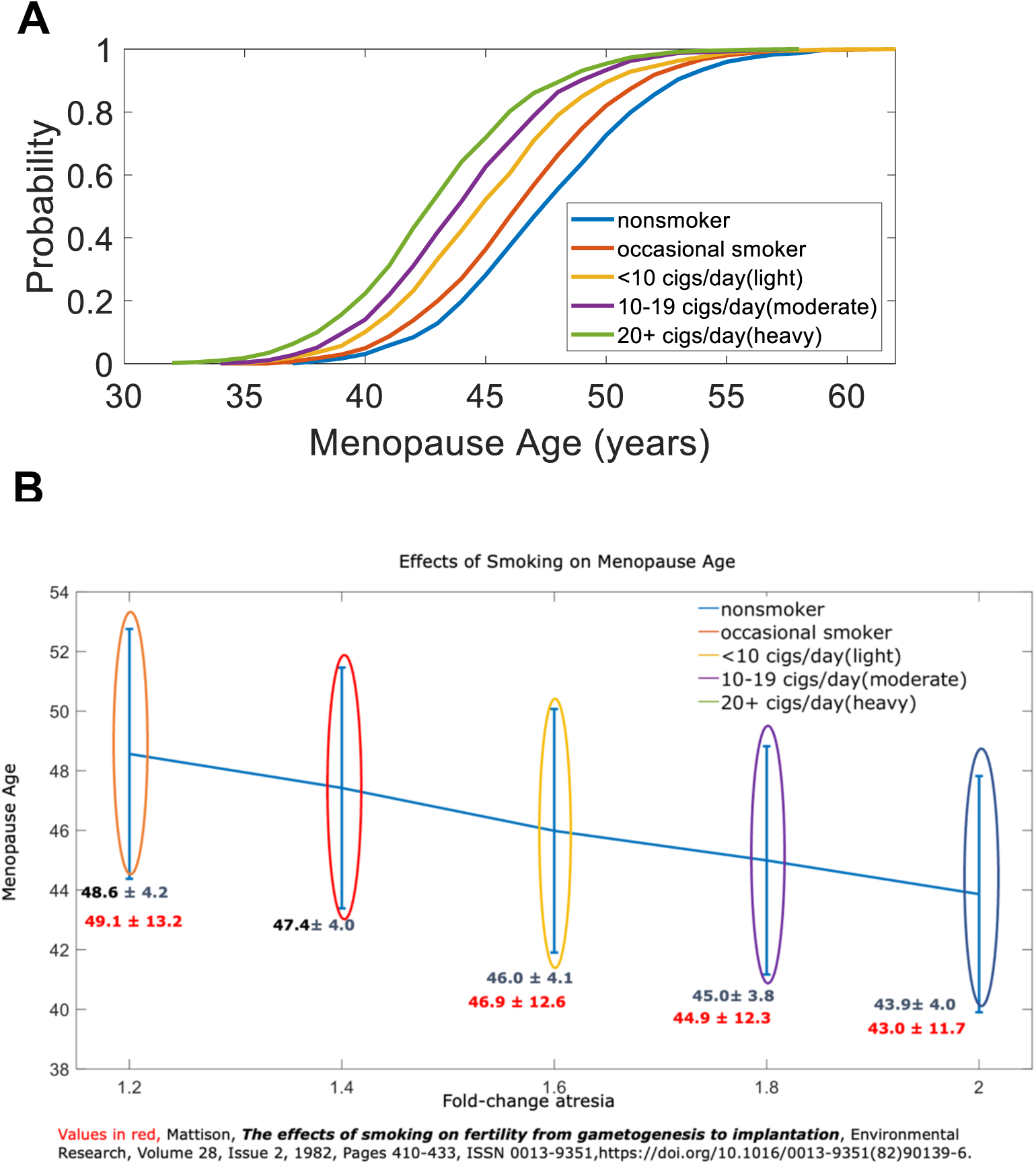
Simulation results for cigarette smoke exposure. **(A)** Cumulative probability of menopause age for different cigarette smoking scenariors as indicated. **(B)** Predicted mean menopause age and sd for different cigarette smoke exposure scenarios as indicated. The simultions were conducted by varying k0 to different fold of the default value from age 15 to 30 as indicated in (B). A 1.2-fold was used to approximate the second-hand smoking scenario for nonsmokers.

## Discussion

In the present study, we constructed a mechanistically based mathematical model that describes ovarian aging. The model clearly recapitulates the biphasic-like depletion behavior of primordial follicles from birth through menopause. It also predicts that the GF depletion rate in the first phase (birth through 30 years) is much smaller than NGF depletion rate, suggesting a relatively constant supply of growing follicles is maintained through most of the reproductively active life stage. The initial endowment or size of the ovarian reserve is the key determinant of menopausal age. When applying the model to exposure scenarios with known or hypothetical mechanisms that deplete NGF and GF, such as chemo-therapy drugs or cigarette smoke, the model recapitulates quantitatively accelerated ovarian aging resulting in earlier predicated age at menopause. To our knowledge, this is the first biologically relevant mechanistic model of follicle depletion for characterizing ovarian aging without relying on empirical mathematical functions.

A prior study by Zhu and colleagues conducted a pooled analysis from 17 observational studies across 7 countries (n = 207,231) to assess the relationship between smoking and age at menopause in the International collaboration for a Life course Approach to reproductive health and Chronic disease Events, or InterLACE (Zhu et al. 2018). They report comparable reductions in age at menopause reported in our study. Compared with 1.7% of never smokers under age 40, 3.1% of current smokers underwent menopause prior to age 40 whereas 43.4% of never smokers report age at menopause of 52+ compared with 29.7% of current smokers. When comparing intensity, they also report a dose dependent trend where higher intensity smoking resulted in more women experiencing menopause earlier. For example, compared with never smokers with the highest proportion of women (>40%) reporting age at menopause age 52 or later, 33.8% of current light smokers report age at menopause 52+, with the majority reporting ages between 45-49 and 50-51 (Zhu et al. 2018). Our model predicts age at menopause 50.7±5.7 years for light smokers. In those reporting moderate (10-19 cigarettes/day) and heavy smoking (20+ cigarettes/day), 33.9 and 35.0%, respectively, report age at menopause of 45-49. Further, moderate and heavy smokers report 27.8% and 26%, respectively, age at menopause 52+, with heavy current smokers reporting the lowest proportion of women experiencing menopause later than age 52 (Zhu et al. 2018). Our model predicts age at menopause 47.3 ± 5.0 years for moderate smokers, and 46.2±4.9 years for heavy smokers.

### Age-dependent primordial follicle fate outcomes

Primordial follicles leave the ovarian reserve pool via either atresia or activation to become growing follicles. Yet, the probabilities of these two fates vary tremendously through the lifetime. NGF are believed to be depleted mainly via atresia in young women but by age 30 and later the recruitment into GF dominates the depletion of NGF (Gougeon 1996). Our simulation result is consistent with this estimate semi-quantitatively. This phenomenon has also been observed in animals. For instance, in mice the primordial follicle activation/atresia probability ratio was estimated to be 0.0036/0.023 before 30 days which then increased to 0.0043/0.0017 after puberty (Faddy, Jones, and Edwards 1976). The fate ratio of 0.024/0.046 per day before 20 days and 0.0034/0.0053 per day after 20 days was also estimated for CD-1 mice (Bristol-Gould et al. 2006), which is approximately equivalent to 155 follicles going through atresia and 81 going to primary follicles per ovary per day before day 20 (Tingen et al. 2009). Therefore, primordial follicle depletion being initially dominated by atresia then by activation appears to be a conserved theme across mammalian species.

### Age-dependent NGF:GF ratio

Also changing with age is the NGF to GF ratio. Our model demonstrates a general declining trend of NGF:GF ratio with age, which has been observed experimentally. In mice, the ratio of primordial to small follicle decreases from about 7:1 to 5:1 to 1:1, from an age of 25 days, 4 months, and 13 months, respectively (Durlinger et al. 1999). In 8-month old mice, the ratio of primordial to primary, secondary, small preantral follicle is slightly higher than 2:1 for each stage, and together primordial to small follicle ratio is 2:3, while the primordial to antral follicle ratio is about 4:1 (Uslu et al. 2017). Although our model correctly recapitulated the downtrend of NGF:GF ratio with age, the absolute value of this ratio appears to be overestimated. Human studies show the primordial to primary follicle ratio can range between 16.5:1 to 41:1 in infants (Forabosco and Sforza 2007), in comparison with the ratio of over 100:1 at a similar age as predicted by our model. This discrepancy can be corrected in future iterations by reducing the atresia rate of GF such that GF can accumulate to high levels. Such revision is not expected to affect the model behaviors qualitatively, as the amount of AMH which is secreted by GF can be easily rescaled.

### Constant supply of GF regulated by intraovarian mechanisms

The most interesting feature predicted by the model is the relatively constant level or supply of GF through most of the reproductive age despite NGF declining by > 90% since birth. This constancy can be explained by feedforward and feedback regulations mediated via InhF and AMH, respectively. Since InhF is proportional to NGF (Eq. 1), NGF cancels out in the 2^nd^ term of Eq. 3 and the activation becomes zero-order at young ages when NGF is high. This explains the relatively constant activation rate and number of GF. Though intraovarian signaling is proportional to the NGF count and density (Schenck et al. 2021), with aging there are fewer NGF which also become sparse, thus the collectively secreted InhF in the local milieu becomes insufficient to keep follicle activation in a zero-order manner by itself. Instead, AMH-mediated feedback starts to predominate to continuously ensure zero-order follicle activation. This feedback mechanism is similar to the end-product exerted feedback in a metabolic pathway that ensures a constant metabolic flux. Its implementation in the setting of ovarian follicle activation provides another example that it is a universal mechanism utilized to achieve constancy or homeostasis.

### Mechanisms and effects of chemo-drug associated menopause

Experimental toxicological studies at human-relevant levels demonstrate Doxorubicin’s ovotoxicity via inhibition of follicle development and oocyte maturation, decreased estradiol secretion, and DNA damage within the granulosa (Xiao et al. 2017), and reduced ovarian size (Ben-Aharon et al. 2010). This is further supported by a systemic review reporting indirect effects, in addition to direct effects, of chemo drugs on POF, namely through impaired somatic cells (i.e. granulosa) and subsequent increased growth activation to replace nonviable follicles (Morgan et al. 2012). A study compared Chinese women undergoing breast cancer chemotherapy (anthracyclines dominant-which usually includes Doxrubicin, or no anthracyclines) aged 40 – 60 years to a control group of women without breast cancer aged 40-65 years. Almost 40% of those undergoing treatment reported being post menopause (mean ages 49.39 ± 5.00 years), compared to 23% controls (mean age 50.53 ± 5.88 years; p<0.001) (Zhao et al. 2024). Another study evaluating age of menopause in pre- or perimenopausal women following treatment for early breast cancer reports median ages across two treatment groups. In women who received one perioperative cycle of cyclophosphamide (CMF), methotrexate, and 5-Fluorouracil (PeCT), or no systemic chemotherapy (no CT), median age of menopause recorded was 50 years (range 34-59), compared with median age 45 years (range 34-55) in those receiving CMF x 6 (repeatedly every 28 days x 6 postoperatively) or CMF x 7 (1 cycle of CMF perioperatively + CMF x 6) (Partridge et al. 2007). A study using menstrual bleeding following breast cancer treatment (varying regimes) as a proxy for ovarian function in women age 20 – 45 years reported menstrual cycles resumed at 6 months post treatment in 85% of women under age 35, and in 61% of women aged 35-40 years (Petrek et al. 2006). This study also reported a reduced odds of menstrual bleeding by 24% for each year age 35 and above (Petrek et al. 2006). Our model of chemo-drug exposure for 1 year at age 20 predicts age of menopause of 47.3 ± 4 years, which is consistent with each of these studies, adding confidence in our model’s performance in predicting age of menopause within a across varying types of treatment protocols and across a range of participants.

### Mechanisms and effects of cigarette smoke toxicity

Cigarette smoke as a mixture is an endocrine disrupting substance, containing ∼7,000 components including hydrocarbons, heavy metals, and aldehydes which have demonstrated associations with diverse reproductive adverse outcomes (Mattison, 1989; Tan et al., 2022). Substantial toxicological data support cigarette smoke induces toxicity on ovarian reserve through direct cytotoxicity via induction of cellular inflammation or oxidative stress, apoptosis due to cellular accumulation of toxicants, and endocrine disruption of hormone signaling (Mattison, 1989). Among which, Benxo(a)Pyrine, (BaP) has exhibited direct effects of toxicity on dormant follicles via DNA damage-induced oxidative stress and lipid peroxidation (Gannon et al., 2012; Kawai, 2012; Siddique et al., 2014; Sobinoff et al., 2012, 2013), granulosa cell autophagy, and accelerated decline of reserve (Gannon 2011, Gannon 2013). Based on these mechanisms, the most straightforward implementation of smoke effect in our model is to increase the atresia rate of the primordial follicles regardless of the specific identity of the active smoke ingredients or their mixtures. Early work (Mattison, 1989) has demonstrated that a 5-20% increase in atresia rate can lead to biologically plausible younger age at menopause. When everything else remains equal, the dose and length of smoking determine the rate of ovarian reserve depletion, and our model predicts the mean menopause age can be accelerated by up to 6-7 years if the smoking history spans teens through early 30s. This model prediction is consistent with findings from early studies (Cooper et al., 1999; Ikuko et al., 1989; Mattison, 1989; Mikkelsen et al., 2007; Whitcomb et al., 2018)

### Limitations and future directions

The size of a woman’s ovary is not constant through her lifetime. Its volume increases nearly 10 fold from 2 through 20 years of age and then declines steadily through menopause, and there appears to be large interindividual variabilities as well (Kelsey et al. 2013). The initial volume growth through puberty is attributed primarily to stromal tissue expansion. This would impact the local concentrations of InhF and AMH, affecting their inhibitory actions on NGF activation, which is not considered in the current model. Therefore, the waning action of InhF with age may be contributed by both a fewer number of NGF and the increasing distance between the follicles. This volume change may also help to explain the observed rising serum AMH from birth through puberty (Kelsey et al. 2011a). In the current model, this time-dependent AMH rise is implemented with a Hill function. It is likely that when the ovary volume is very small, the density of the NGF is so high such that the local InhF concentration is also high, resulting in a strong inhibition of NGF activation. Therefore, the number of GF is quite low at childhood age, manifested as low serum AMH levels. As the ovary volume expands, the local InhF concentration becomes lower, thus lessening the inhibition of NGF activation. As a result, more GF are activated, which then produce higher AMH levels. Therefore, considering age-dependent ovarian volume change in future iterations of the model may help to resolve both issues and underscores the importance of incorporating ovarian spatial information into future model developments. Lastly, AMH plays a role in inhibiting the cyclic recruitment of GF to become large antral follicles to be supplied as waves of follicle cohort in each menstrual cycle (Dumont et al. 2015). This forms another feedforward regulation to further ensure a constant size for each cohort is generated despite variation in the GF pool size. The current model did not consider this route of GF fate regulated by AMH. However, it does not pose an issue in modeling NGF depletion as moving forward to become large antral follicles is only for a small fraction of GF, therefore the overall AMH level will not be significantly affected.

Our findings provide evidence that continued model development holds promise to scalable exploration of mechanisms of follicular depletion and assessment of biologically plausible hypotheses related to perturbations to ovarian reserve that implicate ovarian aging. This model explores dynamics of two biological follicle preservation actions: the primordial follicle pool-originated inhibitory factor and AMH feedback from growing follicles. Although a minimal nonspatial model is presented here, further iterations should incorporate non-random spatial distribution of follicles within the ovary. The non-random distribution may have significant effects on paracrine signaling within the ovary which may have more telling implications of ovarian function given more evidence of follicular clustering with aging (Schenck et al., 2021).

The ideal model for any tool of clinical relevance in predicting reproductive endpoints has important implications for supporting efforts in precision medicine and personalized risk assessment. For instance, accurate predictions of ovarian reserve can help screen for patients who are likely to have poor or hyper-response to ART procedures and prepare the physicians to optimize the dose of gonadotrophins for ovarian stimulation. Implications for a tool as this is especially desirable for predicting menopausal age given current estimated antral follicle count, AMH levels, and other individual-level information characterizing our identified parameters, such as in the cigarette exposure scenario, often have wide interindividual variabilities. This is expected to become even more relevant as we are entering an era of personalized quantification of exposure to EDCs that pose unexplored and unpredictable consequences for fertility such as early natural menopause, chemical-induced genotoxic decay, and other unexpected chemical related perturbations that accelerate ovarian aging thereby potentially decreasing actualized reproductive years.

A model for age-related follicular depletion, menopausal age, as well as early menopausal age should be integrated with models that will also predict complex reproductive outcomes such as fertility and fecundity. Fertility and fecundity may be polygenic and sensitive to epigenetic markers therefore, predictive models should ideally include both qualitative and quantitative measures of oocyte, follicular, and reproductive health which may be extracted from health records and studies evaluating population-level markers. Integrating multiple models developed for different but related purposes in female reproduction may lead to the development of digital twins of women (Katsoulakis et al. 2024).

## Conclusions

Our model recapitulates the biphasic depletion kinetic of NGF and GF from birth through menopause and accurately predicts menopausal age distribution in humans. Further, the model predicts a relatively constant supply of growing follicles through most of the reproductively active life stage. The initial size of the ovarian reserve is the key determinant of the menopausal age so maximizing the ovarian reserve at birth and preserving this supply across the life course the key parameters to ensuring a long reproductive life span, including an older age at menopause. This model predicts mean menopause age ± SD of 50.3 ± 4.3 years, which is comparable with national and global estimates of reproductive senescence. Further, the model demonstrates a dose-dependent reduction in menopause age with increased exposure to tobacco smoking. This contributes to the extensive body of evidence of cigarette smoke-induced reproductive toxicity via reduction in years of reproductive life and earlier age at menopause. This model holds promise in predicting chemically induced POI and early menopause, indicating potential utility in modelling the possible downstream effects of quantifiable chemical exposures that act upon pathways involved in ovarian regulation.

## References

Anckaert, E., J. Smitz, J. Schiettecatte, B. M. Klein, and J. C. Arce. 2012. ‘The value of anti-Mullerian hormone measurement in the long GnRH agonist protocol: association with ovarian response and gonadotrophin-dose adjustments’, Hum Reprod, 27: 1829–39.

Ben-Aharon, I., H. Bar-Joseph, G. Tzarfaty, L. Kuchinsky, S. Rizel, S. M. Stemmer, and R. Shalgi. 2010. ‘Doxorubicin-induced ovarian toxicity’, Reproductive biology and endocrinology : RB&E, 8: 20.

Bristol-Gould, S. K., P. K. Kreeger, C. G. Selkirk, S. M. Kilen, K. E. Mayo, L. D. Shea, and T. K. Woodruff. 2006. ‘Fate of the initial follicle pool: empirical and mathematical evidence supporting its sufficiency for adult fertility’, Dev Biol, 298: 149–54.

Broekmans, F. J., M. R. Soules, and B. C. Fauser. 2009. ‘Ovarian aging: mechanisms and clinical consequences’, Endocr Rev, 30: 465–93.

Da Silva-Buttkus, P., G. Marcelli, S. Franks, J. Stark, and K. Hardy. 2009. ‘Inferring biological mechanisms from spatial analysis: prediction of a local inhibitor in the ovary’, Proc Natl Acad Sci U S A, 106: 456–61.

Ding, N., S. D. Harlow, J. F. Randolph, Jr., R. Loch-Caruso, and S. K. Park. 2020. ‘Perfluoroalkyl and polyfluoroalkyl substances (PFAS) and their effects on the ovary’, Hum Reprod Update, 26: 724–52.

’<dissertation_dingning_1.pdf>’.

Dumont, Agathe, Geoffroy Robin, Sophie Catteau-Jonard, and Didier Dewailly. 2015. ‘Role of Anti-Müllerian Hormone in pathophysiology, diagnosis and treatment of Polycystic Ovary Syndrome: a review’, Reproductive Biology and Endocrinology, 13: 137.

Durlinger, A. L., P. Kramer, B. Karels, F. H. de Jong, J. T. Uilenbroek, J. A. Grootegoed, and A. P. Themmen. 1999. ‘Control of primordial follicle recruitment by anti-Müllerian hormone in the mouse ovary’, Endocrinology, 140: 5789–96.

Faddy, M. J., and R. G. Gosden. 1996. ‘A model conforming the decline in follicle numbers to the age of menopause in women’, Hum Reprod, 11: 1484–6.

Faddy, M. J., R. G. Gosden, A. Gougeon, S. J. Richardson, and J. F. Nelson. 1992. ‘Accelerated disappearance of ovarian follicles in mid-life: implications for forecasting menopause’, Hum Reprod, 7: 1342–6.

Faddy, M. J., E. C. Jones, and R. G. Edwards. 1976. ‘An analytical model for ovarian follicle dynamics’, J Exp Zool, 197: 173–85.

Forabosco, A., and C. Sforza. 2007. ‘Establishment of ovarian reserve: a quantitative morphometric study of the developing human ovary’, Fertil Steril, 88: 675–83.

Gougeon, A. 1996. ‘Regulation of ovarian follicular development in primates: facts and hypotheses’, Endocr Rev, 17: 121–55.

Gougeon, A., R. Ecochard, and J. C. Thalabard. 1994. ‘Age-related changes of the population of human ovarian follicles: increase in the disappearance rate of non-growing and early-growing follicles in aging women’, Biol Reprod, 50: 653–63.

Gynecologists, American College of Obstetricians and. 2014. “Primary ovarian insufficiency in adolescents and young women.” In Committee Opinion. Obstet Gynecol.

Hagen, C. P., L. Aksglaede, K. Sørensen, K. M. Main, M. Boas, L. Cleemann, K. Holm, C. H. Gravholt, A. M. Andersson, A. T. Pedersen, J. H. Petersen, A. Linneberg, S. Kjaergaard, and A. Juul. 2010. ‘Serum levels of anti-Müllerian hormone as a marker of ovarian function in 926 healthy females from birth to adulthood and in 172 Turner syndrome patients’, J Clin Endocrinol Metab, 95: 5003–10.

Hansen, K. R., N. S. Knowlton, A. C. Thyer, J. S. Charleston, M. R. Soules, and N. A. Klein. 2008. ‘A new model of reproductive aging: the decline in ovarian non-growing follicle number from birth to menopause’, Hum Reprod, 23: 699–708.

Hirshfield, A. 1994a. ’Primordial follicles inhibits onset of follicle growth’. Hirshfield, A. N. 1994b. ‘Relationship between the supply of primordial follicles and the onset of follicular growth in rats’, Biol Reprod, 50: 421–8.

Iyer, Sandhya, K Kallathikumar, Prachi Sinkar, and Amruta Velumani. 2019. ‘Anti-Mullerian hormone (AMH) and Age–An Indian laboratory retrospective analysis’, Asian Journal of Health Sciences, 5: 7–7.

Kalich-Philosoph, L., H. Roness, A. Carmely, M. Fishel-Bartal, H. Ligumsky, S. Paglin, I. Wolf, H. Kanety, B. Sredni, and D. Meirow. 2013. ‘Cyclophosphamide triggers follicle activation and “burnout”; AS101 prevents follicle loss and preserves fertility’, Sci Transl Med, 5: 185ra62.

Katsoulakis, Evangelia, Qi Wang, Huanmei Wu, Leili Shahriyari, Richard Fletcher, Jinwei Liu, Luke Achenie, Hongfang Liu, Pamela Jackson, Ying Xiao, Tanveer Syeda-Mahmood, Richard Tuli, and Jun Deng. 2024. ‘Digital twins for health: a scoping review’, npj Digital Medicine, 7: 77.

Kelsey, T. W., S. K. Dodwell, A. G. Wilkinson, T. Greve, C. Y. Andersen, R. A. Anderson, and W. H. Wallace. 2013. ‘Ovarian volume throughout life: a validated normative model’, PLoS One, 8: e71465.

Kelsey, T. W., P. Wright, S. M. Nelson, R. A. Anderson, and W. H. Wallace. 2011a. ‘A validated model of serum anti-müllerian hormone from conception to menopause’, PloS one, 6: e22024.

Kelsey, Thomas W., Phoebe Wright, Scott M. Nelson, Richard A. Anderson, and W. Hamish B. Wallace. 2011b. ‘A Validated Model of Serum Anti-Müllerian Hormone from Conception to Menopause’, PLoS One, 6: e22024.

Lie Fong, S., J. A. Visser, C. K. Welt, Y. B. de Rijke, M. J. Eijkemans, F. J. Broekmans, E. M. Roes, W. H. Peters, A. C. Hokken-Koelega, B. C. Fauser, A. P. Themmen, F. H. de Jong, I. Schipper, and J. S. Laven. 2012. ‘Serum anti-müllerian hormone levels in healthy females: a nomogram ranging from infancy to adulthood’, J Clin Endocrinol Metab, 97: 4650–5.

Messerlian, Carmen, Paige L. Williams, Jennifer B. Ford, Jorge E. Chavarro, Lidia Mínguez-Alarcón, Ramace Dadd, Joseph M. Braun, Audrey J. Gaskins, John D. Meeker, Tamarra James-Todd, Yu-Han Chiu, Feiby L. Nassan, Irene Souter, John Petrozza, Myra Keller, Thomas L. Toth, Antonia M. Calafat, Russ Hauser, and Earth Study Team for the. 2018. ‘The Environment and Reproductive Health (EARTH) Study: a prospective preconception cohort’, *Human Reproduction Open*, 2018.

Morgan, S., R. A. Anderson, C. Gourley, W. H. Wallace, and N. Spears. 2012. ‘How do chemotherapeutic agents damage the ovary?’, Hum Reprod Update, 18: 525–35.

Partridge, Ann, Shari Gelber, Richard D. Gelber, Monica Castiglione-Gertsch, Aron Goldhirsch, and Eric Winer. 2007. ‘Age of menopause among women who remain premenopausal following treatment for early breast cancer: Long-term results from International Breast Cancer Study Group Trials V and VI’, European Journal of Cancer, 43: 1646–53.

Pastore, L. M., M. S. Christianson, J. Stelling, W. G. Kearns, and J. H. Segars. 2018. ‘Reproductive ovarian testing and the alphabet soup of diagnoses: DOR, POI, POF, POR, and FOR’, J Assist Reprod Genet, 35: 17–23.

Petrek, Jeanne A., Michelle J. Naughton, L. Douglas Case, Electra D. Paskett, Elizabeth Z. Naftalis, S. Eva Singletary, and Paniti Sukumvanich. 2006. ‘Incidence, Time Course, and Determinants of Menstrual Bleeding After Breast Cancer Treatment: A Prospective Study’, Journal of Clinical Oncology, 24: 1045–51.

Podfigurna-Stopa, A., A. Czyzyk, M. Grymowicz, R. Smolarczyk, K. Katulski, K. Czajkowski, and B. Meczekalski. 2016. ‘Premature ovarian insufficiency: the context of long-term effects’, J Endocrinol Invest, 39: 983–90.

Promotion, National Center for Chronic Disease Prevention and Health. 2022. “Infertility FAQs.” In, edited by National Center for Chronic Disease Prevention and Health Promotion Division of Reproductive Health.

Schenck, A., M. Vera-Rodriguez, G. Greggains, B. Davidson, and P. Fedorcsák. 2021. ‘Spatial and temporal changes in follicle distribution in the human ovarian cortex’, Reprod Biomed Online, 42: 375–83.

Seifer, D. B., V. L. Baker, and B. Leader. 2011. ‘Age-specific serum anti-Müllerian hormone values for 17,120 women presenting to fertility centers within the United States’, Fertil Steril, 95: 747–50.

Tingen, C. M., S. K. Bristol-Gould, S. E. Kiesewetter, J. T. Wellington, L. Shea, and T. K. Woodruff. 2009. ‘Prepubertal primordial follicle loss in mice is not due to classical apoptotic pathways’, Biol Reprod, 81: 16–25.

Uslu, B., C. C. Dioguardi, M. Haynes, D. Q. Miao, M. Kurus, G. Hoffman, and J. Johnson. 2017. ‘Quantifying growing versus non-growing ovarian follicles in the mouse’, J Ovarian Res, 10: 3.

Wallace, W. H., and T. W. Kelsey. 2010. ‘Human ovarian reserve from conception to the menopause’, PLoS One, 5: e8772.

Wang, Y., M. Liu, S. B. Johnson, G. Yuan, A. K. Arriba, M. E. Zubizarreta, S. Chatterjee, M. Nagarkatti, P. Nagarkatti, and S. Xiao. 2019a. ‘Doxorubicin obliterates mouse ovarian reserve through both primordial follicle atresia and overactivation’, Toxicol Appl Pharmacol, 381: 114714.

Wang, Y., M. Liu, S. B. Johnson, G. Yuan, A. K. Arriba, M. E. Zubizarreta, S. Chatterjee, M. Nagarkatti, P. Nagarkatti, and S. Xiao. 2019b. ‘Doxorubicin obliterates mouse ovarian reserve through both primordial follicle atresia and overactivation’, Toxicol Appl Pharmacol: 114714.

Xiao, S., J. Zhang, M. Liu, H. Iwahata, H. B. Rogers, and T. K. Woodruff. 2017. ‘Doxorubicin Has Dose-Dependent Toxicity on Mouse Ovarian Follicle Development, Hormone Secretion, and Oocyte Maturation’, Toxicological sciences : an official journal of the Society of Toxicology, 157: 320–29.

Xu, J., Y. Wang, A. E. Kauffman, Y. Zhang, Y. Li, J. Zhu, K. Maratea, K. Fabre, Q. Zhang, T. K. Woodruff, and S. Xiao. 2020. ‘A Tiered Female Ovarian Toxicity Screening Identifies Toxic Effects of Checkpoint Kinase 1 Inhibitors on Murine Growing Follicles’, Toxicol Sci, 177: 405–19.

Zeleznik, Anthony. 2004. ’The physiology of follicle selection’.

Zhao, D., Q. Ma, G. Li, R. Qin, Y. Meng, and P. Li. 2024. ‘Treatment-induced menopause symptoms among women with breast cancer undergoing chemotherapy in China: a comparison to age- and menopause status-matched controls’, Menopause, 31: 145–53.

Zhao, Y., H. Feng, Y. Zhang, J. V. Zhang, X. Wang, D. Liu, T. Wang, R. H. W. Li, E. H. Y. Ng, W. S. B. Yeung, K. A. Rodriguez-Wallberg, and K. Liu. 2021. ‘Current Understandings of Core Pathways for the Activation of Mammalian Primordial Follicles’, Cells, 10.

Zhu, D., H. F. Chung, N. Pandeya, A. J. Dobson, J. E. Cade, D. C. Greenwood, S. L. Crawford, N. E. Avis, E. B. Gold, E. S. Mitchell, N. F. Woods, D. Anderson, D. E. Brown, L. L. Sievert, E. J. Brunner, D. Kuh, R. Hardy, K. Hayashi, J. S. Lee, H. Mizunuma, G. G. Giles, F. Bruinsma, T. Tillin, M. K. Simonsen, H. O. Adami, E. Weiderpass, M. Canonico, M. L. Ancelin, P. Demakakos, and G. D. Mishra. 2018. ‘Relationships between intensity, duration, cumulative dose, and timing of smoking with age at menopause: A pooled analysis of individual data from 17 observational studies’, PLoS Med, 15: e1002704.

